# Acquired loss of cardiac vagal activity is associated with myocardial injury in patients undergoing non-cardiac surgery: prospective observational mechanistic cohort study

**DOI:** 10.1101/623165

**Authors:** Shaun M. May, Anna Reyes, Gladys Martir, Joseph Reynolds, Laura Gallego Paredes, Shamir Karmali, Robert CM Stephens, David Brealey, Gareth L. Ackland

## Abstract

**Background:** Myocardial injury is more frequent after non-cardiac surgery in patients with preoperative cardiac vagal dysfunction as quantified by delayed heart rate recovery after cessation of cardiopulmonary exercise testing. Here, we hypothesised that serial and dynamic measures of perioperative cardiac vagal activity should also be associated with myocardial injury after non-cardiac surgery.

**Methods:** Serial measures in cardiac vagal activity were quantified preoperatively and daily using heart rate variability and a standardised orthostatic challenge in patients undergoing elective non-cardiac surgery. The primary outcome was myocardial injury (high-sensitivity troponin (hsTnT) ≥15ng.L^−1^) within 48h of surgery. Clinicians, patients and investigators were blinded to hsTnT. The exposure of interest was cardiac vagal activity (high-frequency power spectral analysis [HF_log_]) and heart rate recovery after a standardised orthostatic challenge.

**Results:** hsTnT≥15ng.L^−1^ occurred in 48/189 [25%] patients, of whom 41/48 [85%] had a revised cardiac risk index score <2. Patients with a post surgery troponin HsTnT ≥15ng.L^−1^ were associated with an early loss (within 24h) of cardiac vagal activity (HF_Log_) post surgery compared to day of surgery (4.19 [95%CI:3.62-4.75] vs 5.22 [95%CI:4.64-5.81]; p<0.001). Heart rate recovery after a standardised orthostatic challenge after surgery was slower in patients with hsTnT≥15ng.L^−1^ (5 beats minute^−1^ (95% CI: 3 −7), compared to heart rate recovery in patients who remained free of myocardial injury (10 beats minute^−1^ (95%CI:7 to 12]; p = 0.02).

**Conclusions:** Real-time, serial heart rate measures indicating loss of cardiac vagal activity are associated with perioperative myocardial injury in lower-risk patients undergoing non-cardiac surgery.

## Introduction

Perioperative myocardial injury (PMI) characterised by high-sensitivity troponin >99^th^ centile after non-cardiac surgery, occurs ∼25% patients (1) and is associated with increased risk of complications including death within 30 days of surgery (2). At least 70% of non-cardiac surgical patients with raised post-surgery troponin levels demonstrate no ischaemic features (1, 2), suggesting mechanisms other than ischaemia or thrombosis may be involved.

In the Measurement of exercise Tolerance before Surgery (METS) study, delayed heart rate recovery (HRR) quantified after cessation of preoperative cardiopulmonary exercise testing, the gold-standard measure of cardiac vagal activity(3), was the sole physiological parameter associated with post-surgery myocardial injury. (4) Delayed HRR is also strongly associated with baroreflex dysfunction (5) which predisposes patients to haemodynamic compromise linked to post operative morbidity (6). Patients with impaired cardiac vagal activity acquired during surgery may, therefore, be at particular risk of sustaining PMI after major non-cardiac surgery. A readily testable measure of cardiac vagal function after orthostatic challenge is heart rate recovery after standardised orthostatic challenge (7, 8).

Since our previous approach only defined pre-surgery cardiac vagal impairment, we sought to examine whether serial static and dynamic measures of cardiac (vagal) dysfunction, may also be associated with perioperative myocardial injury. We employed heart rate variability and a standardised orthostatic challenge to quantify, in real-time, changes in cardiac vagal activity throughout the perioperative period.

## Methods

### Study design

This was a single centre, prospective observational mechanistic cohort study conducted at University College London Hospital from 05/10/2016 to 14/01/2019. The study was approved by a research ethics committee and conducted in accordance with the principles of the Declaration of Helsinki and the Research Governance Framework. (MREC: 16/LO/06/35). The Strengthening and Reporting of observational Studies in Epidemiology (STROBE) guideline(9) were followed and a pre-publication analysis plan has been available at www.ucl.ac.uk/anaesthesia/trials (published online 01/12/2018). Written informed consent was obtained from all patients before surgery.

### Inclusion and exclusion criteria

Adult patients undergoing major elective surgery with a planned overnight stay in hospital were eligible for recruitment provided they satisfied the following criteria: American Society Anaesthesiology score ≥3, major surgery with expected surgical operating time ≥ 120 minutes. Exclusion criteria were current atrial and/or ventricular arrhythmia, (including permanent or temporary pacemaker in-situ) and refusal of consent.

### Primary outcome

The primary outcome was perioperative myocardial injury (PMI), defined as high sensitivity troponin (HsTnT) ≥15ng.L^−1^ within two days of surgery. The cut off value is above the limit for the normal range for the Troponin-T high sensitivity Cobas assay (10). Plasma troponin levels were batch analysed after the patients left hospital by an independent laboratory blinded to patient details (The Doctors Laboratory, United Kingdom).

### Secondary outcomes

The secondary outcome was all-cause postoperative morbidity within 5 days after surgery, assessed using the Post Operative Morbidity Survey(11) (POMS; Supplementary Table 1), which was collected prospectively on post-surgery day 3 and 5. The time to become morbidity-free and length of hospital stay were also calculated.

**Table 1:** Baseline patient characteristics. Data is presented as mean with standard deviations (SD) for parametric data and as median (25th-75th interquartile range) for non-parametric data. Frequencies are presented with percentages (%). Age is rounded to the nearest year. PMI: perioperative myocardial injury (HS TnT ≤14 ng/L). CKD: chronic kidney disease. Stage is defined as per KDIGO recommendations (29). RCRI: Revised cardiac risk index score. ACE-I: Angiotensin converting enzyme inhibitor. ARB: angiotensin receptor blocker. Intraoperative hypotension is defined as ≥1 intraoperative episode after induction of anaesthesia of systolic blood pressure <90mmHg. Other units as indicated. Statistical analysis using Student’s unpaired t-test for continuous data and the χ2 test for categorical data.

|  | Whole cohort<br>N=189 | hsTnT ≤14ng L<sup>-1</sup><br>n= 141 | hsTnT ≥15ng L<sup>-1</sup><br>n= 48 | p-value |
| --- | --- | --- | --- | --- |
| Post operation Troponin Peak ng L<sup>-1</sup> | 9 (15-22) | 7 (5-11) | 21 (17-23) | <0.001 |
| **Clinical Characteristics** |  |  |  |  |
| Age (yr) | 67 (60-76) | 66 (58-72) | 74 (66-78) | <0.001 |
| Male sex (n;%) | 81 (42.9%) | 52 (36.9%) | 29 (60.4%) | 0.004 |
| Body Mass Index (kg m<sup>-2</sup>) | 27.5 (24.4-31.3) | 28.0(24.6-31.4) | 26.1 (24.1-31.2) | 0.131 |
| CKD Stage ≥2 | 110 (58.2%) | 80 (57.6%) | 30 (62.5%) | 0.485 |
| Haemoglobin (g L<sup>-1</sup>) | 13.1 (11.7-14.1) | 13.2 (12.2-14.2) | 12.2 (11.1-13.8) | 0.054 |
| **Co-morbidities (n; %)** |  |  |  |  |
| RCRI ≥2 | 15 (7.9%) | 8 (5.8%) | 7 (14.6%) | 0.049 |
| Chronic obstructive airway disease | 15 (7.9%) | 12 (8.5%) | 3 (6.3%) | 0.617 |
| Asthma | 29 (15.3%) | 21 (14.9%) | 8 (16.7%) | 0.769 |
| Ischaemic heart disease | 11 (5.8%) | 6 (4.3%) | 5 (10.4%) | 0.115 |
| Diabetes mellitus | 41 (21.7%) | 32(22.7%) | 9 (18.8%) | 0.567 |
| Congestive heart failure | 2 (1.1%) | 1 (0.7%) | 1 (2.1%) | 0.422 |
| Cerebral vascular disease | 11 (5.8%) | 6 (4.3%) | 5 (10.4%) | 0.115 |
| Peripheral vascular disease | 7 (3.7%) | 5 (3.6%) | 2 (4.2%) | 0.844 |
| Hypertension | 104 (55.0%) | 74 (52.5%) | 30 (62.5%) | 0.228 |
| Active cancer | 106 (56.1%) | 73 (51.8%) | 33 (68.8%) | 0.041 |
| **At least one cardiovascular medication (n; %)** | **92 (48.7%)** | **66 (46.8%)** | **26 (54.2%)** | **0.378** |
| Beta blocker | 21 (11.1%) | 12 (8.5%) | 9 (18.8%) | 0.051 |
| Calcium channel blocker | 49 (25.9%) | 35 (24.82%) | 14 (29.2%) | 0.553 |
| Doxazosin | 9 (4.8%) | 5 (3.6%) | 4 (8.3%) | 0.179 |
| **ARB/ACE-I** | **53 (28.0%)** | **37 (26.2%)** | **16 (33.3%)** | **0.345** |
| **Type of surgery (n; %)** |  |  |  |  |
| Gastrointestinal | 95 (50.3%) | 66 (46.8%) | 29 (60.4%) | 0.103 |
| Urology/gynaecological | 51 (29.9%) | 41 (29.1%) | 10 (20.8%) | 0.266 |
| Head/neck/ear, nose, throat | 2 (1.1%) | 2 (1.4%) | 0 (0.0%) | 0.407 |
| Orthopaedic | 41 (21.7%) | 32 (22.7%) | 9 (18.8%) | 0.567 |
| **Type of anaesthesia** |  |  |  |  |
| General | 56 (29.6%) | 47 (33.3%) | 9 (18.8 %) | 0.056 |
| General + epidural | 90 (47.6%) | 62 (44.0%) | 28 (58.3%) | 0.085 |
| General anaesthesia +spinal | 30 (15.9%) | 24 (17.0%) | 6 (12.5%) | 0.717 |
| Neuroaxial only | 13 (6.9%) | 8 (5.7%) | 5 (10.4%) | 0.154 |
| **Intraoperative events** |  |  |  |  |
| Duration of surgery (minutes) | 296 (135-300) | 184 (129-260) | 255 (169-383) | 0.001 |
| Intraoperative hypotension (n,%) | 114 (60.3%) | 84 (59.6%) | 30 (62.5%) | 0.721 |

### Explanatory variables - Cardiac autonomic activity

Lifecard CF digital Holter monitors (Spacelabs Healthcare, Hertford UK) were used to capture three lead electrocardiogram (ECG) recordings. The ECGs were analysed on Kubios HRV Premium software Version 3.1.0 (Kubios HRV 2017, Finland)(12). Holter data cleaning and analysis were undertaken blinded to perioperative data and outcomes. Heart rate variability analysis was calculated for 5 minute segments according to Taskforce guidelines(13). (Detailed methodology is provided in the Supplementary data.). Serial changes in cardiac autonomic activity were quantified using three established measures of autonomic modulation of heart rate; time domain, frequency domain and orthostatic modulation of heart rate.

### Time domain measures

We used the square root of the mean of the sum of the squares of the successive differences between adjacent beat-to beat intervals (RMSSD) and Standard Deviation of the N-N interval (SDNN), which both reflect cardiac parasympathetic activity(13)

### Frequency domain measures

We used high frequency (0.15-0.4Hz) power spectral analysis of R-R interval time series as a measure of cardiac parasympathetic activity(13). Low frequency (0.04-0.15Hz) power spectral analysis was also assessed as a measure of arterial baroreflex activity (14).

### Orthostatic autonomic changes

Heart rate was measured continuously during an orthostatic challenge. Average heart rate at 10 second intervals was calculated. The 10 second bin interval where peak heart rate occurred after the orthostatic challenge was identified for each individual patient. The change in heart rate (heart rate recovery) after peak heart rate was calculated for each patient at each 10s interval after time peak heart rate was identified. Detailed methodology is provided in Supplementary data.

### Experimental protocol

The protocol for the study is summarised in (FIGURE 1). On the morning of surgery continuous three lead electrocardiogram recordings were made in a quiet clinical environment. After 10 minutes recording in the supine position, patients were given an orthostatic challenge by elevating to a 45 degrees head-up position using an electronically adjustable hospital bed (Model 2232, ArjoHuntleigh, Sweden). A further 10 min recording was made in the 45 degree head-up position. Blood pressure and respiratory rate observations were made at the beginning of each 10 min recording. The protocol was then repeated 24h and 48h after surgery.

**Figure 1:**
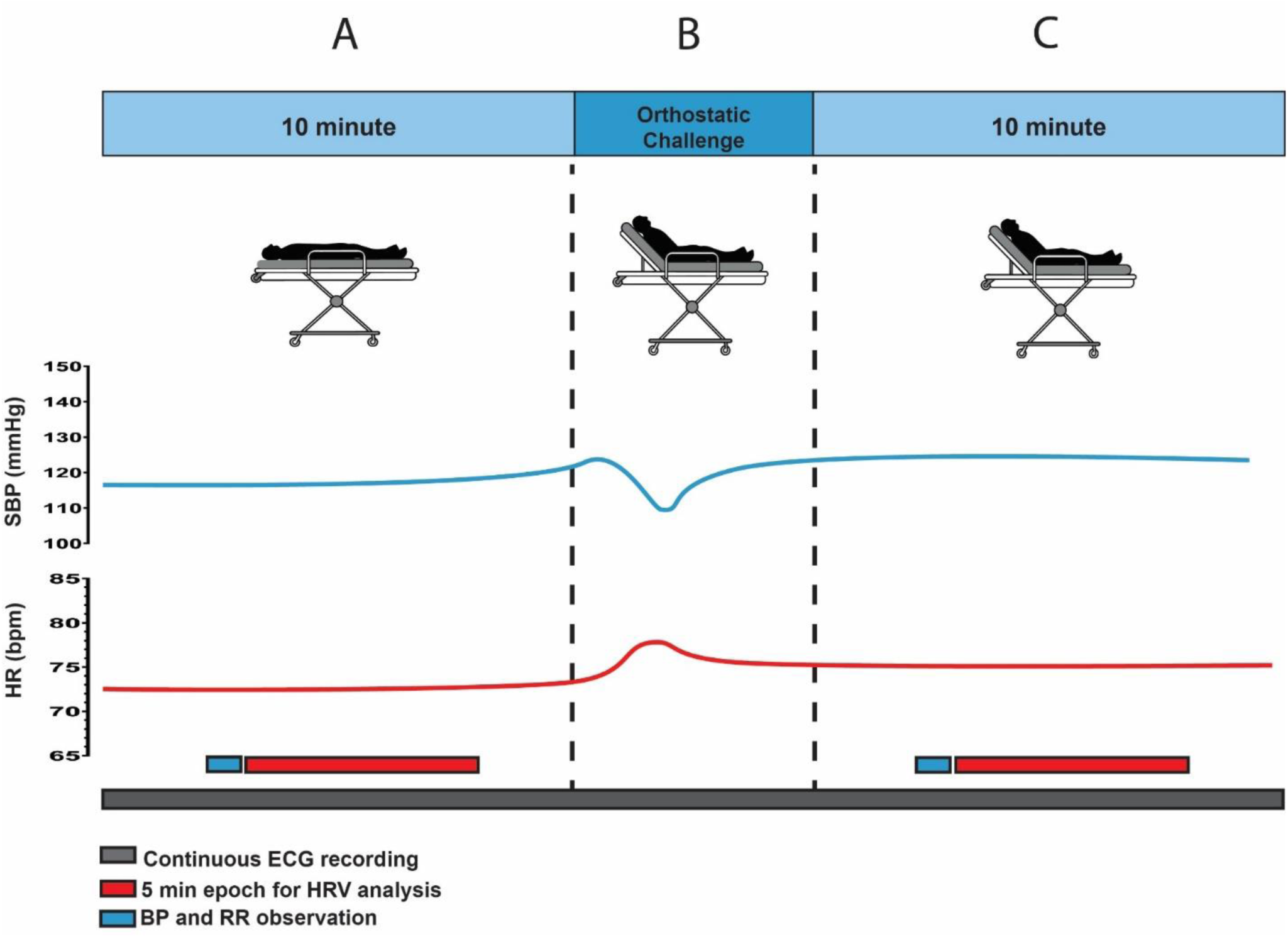
Experimental protocol: A: 10 minute supine position. B:Transition from supine to 45 degree head up position (Orthostatic challenge). C: 10 minute 45 degree head up position. ECG: electrocardiogram. BP: blood pressure. RR: respiratory rate. Heart rate (HR)-red and Systolic blood pressure (SBP) blue predicted response with an orthostatic challenge.

### Perioperative management

Surgery and anaesthesia were conducted by consultant staff and intraoperative care was delivered according to usual standard of care. Postoperative management was conducted according to local enhanced recovery after surgery clinical guidelines. All clinicians were blinded to holter data and troponin results.

### Statistical analysis

Manual and automated validation checks of data were undertaken. Categorical data are summarised as absolute values (percentage). Continuous data are presented as mean (SD), unless stated otherwise. We present participants’ characteristics stratified by PMI outcome. Continuous longitudinal data were analysed using a three-way Mixed Model repeated measures analysis. The fixed effects were; PMI (HsTnT ≥15ng L^−1^ and HsTnT ≤14ng L^−1^), day relating to surgery (day of surgery, 24h after surgery and 48h after surgery) and body position (supine and 45 degrees sitting). Individual comparisons between groups were calculated using post-hoc Bonferroni tests. Associations with perioperative myocardial injury were assessed by logistic regression analysis, taking into account age, gender, Revised Cardiac Risk Index and HRV measures of cardiac parasympathetic activity. The HRV variables were ranked and divided into tertiles and each tertile treated as a categorical value. Length of hospital stay and time to become morbidity free were analysed using Kaplan-Meier Analysis. P<0.05 was considered statistically significant. All statistical analyses were undertaken using NCSS 12 (Kaysville, UT, USA).

### Sample size calculation

Our previous work has demonstrated that cardiac vagal dysfunction as defined by abnormal heart rate recovery ≤12 beats minute^−1^ is present in ∼35% surgical patients (15). Perioperative myocardial injury occurs in ∼25% non cardiac surgical patient(2). In METS, we observed a 50% increase in myocardial injury in patients with preoperative vagal dysfunction (OR, 1.54 (1.11-2.13); P = 0.009). (4) Therefore a sample size of for the study would be at least 152 patients (alpha = 0.05 and beta = 0.9) (16). Allowing for ∼15% dropout, and/or cancellation of surgery and ∼25% incomplete ECG data (17) we estimated that at least 210 patients would be required to demonstrate to detect differences in ECG-derived measures of parasympathetic dysfunction.

## Results

210 patients were recruited into the study between October 2016 and December 2018. After predefined exclusions, we analysed data obtained from 189 participants (Figure 2). The proportion of patient with a RCRI < 2 was 92.1% (174/189).

**Figure 2:**
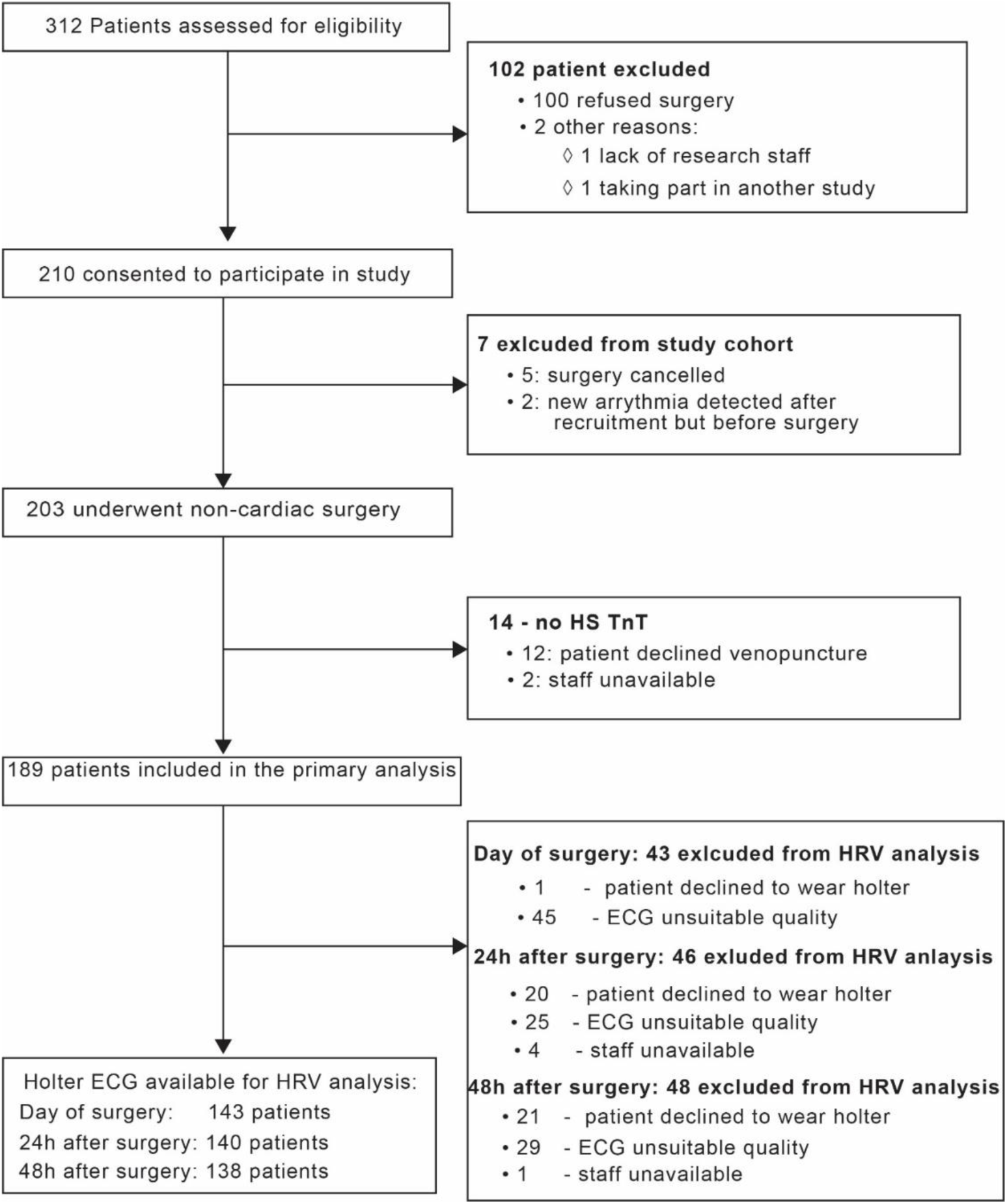
Consort diagram showing patients included in the analysis. HS TnT: High sensitivy Troponin T assay

### Primary outcome: perioperative myocardial injury

48/189 (25%) patients sustained PMI within 2 days after surgery. Median troponin values were 21ng L^−1^ (17-23ng L^−1^) in patients with PMI, compared to patients who failed to breach the 99^th^ centile value, 7 ng L^−1^ (5-11 ng L^−1^), p <0.001). Resting supine heart rate, respiratory rate, systolic and diastolic blood pressure before surgery were similar in both groups (Figure 3).

**Figure 3:**
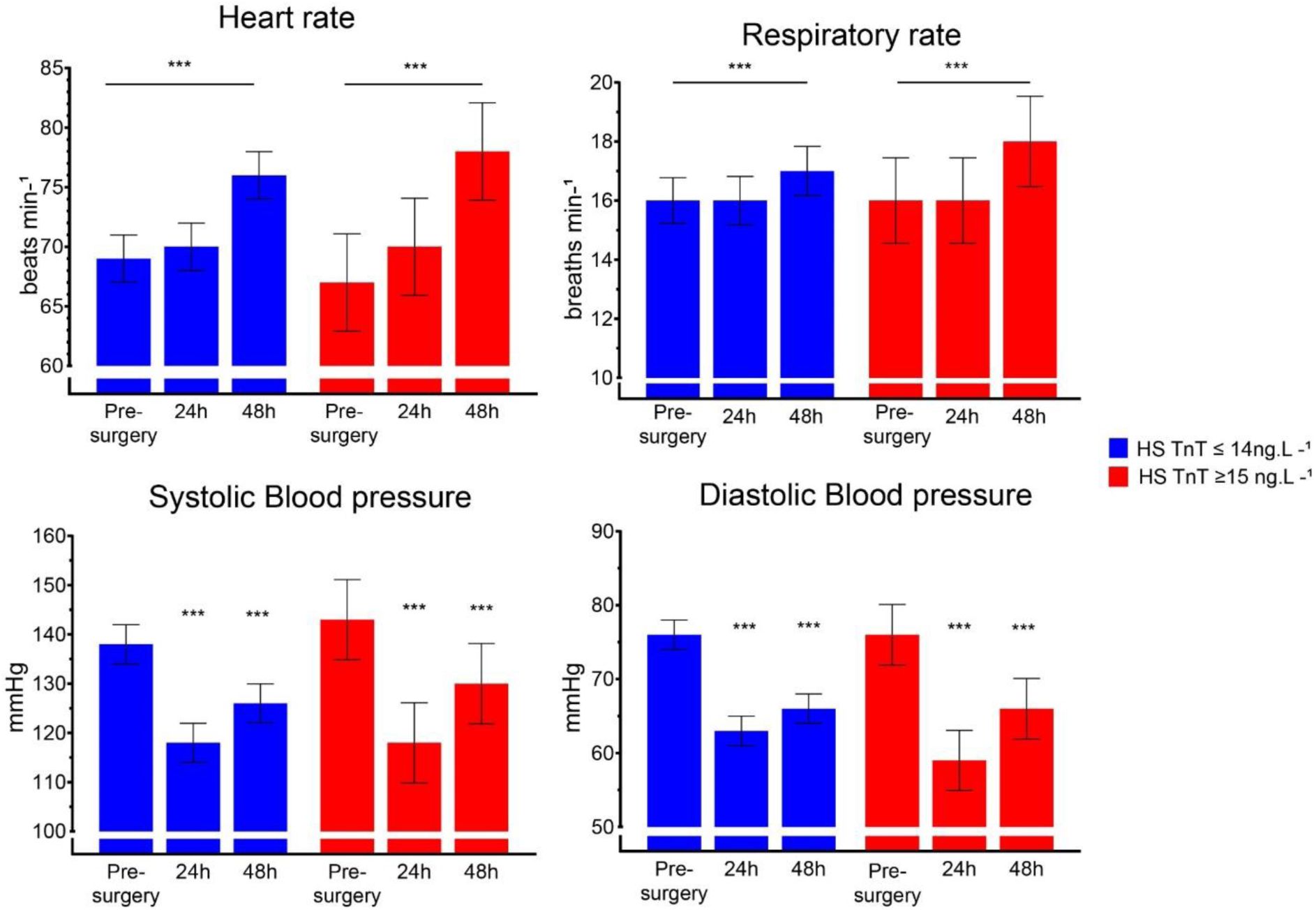
Serial resting supine haemodynamic and respiratory variables. Data is presented as mean values with 95% CI. P value refers to Pre-surgery vs 24h after surgery and Pre-surgery vs 48h after surgery (analysed by Mixed model repeated measures, Bonferroni’s Test). ** p<0.01, *** p<0.001

### Secondary outcome: postoperative morbidity

Patients who developed PMI after surgery had a higher proportion of POMS, 41/48 (85.4%) versus 91/141 (64.5%) who did not develop PMI; (OR: 3.41(95% CI:1.29-9.09); p=0.007). In addition those with raised troponin post surgery had a higher proportion of cardiovascular associated POMS within 5 days after surgery (27/48 (56.3%) vs 29/141 (20.5%),OR 4.97 (95% CI: 1.48-8.45); p <0.001. However no patient had a documented myocardial infarction after surgery. Individualised morbidity data for postoperative days 3 and 5 and length of stay data are provided in Supplementary Data Table 2.

### Explanatory variables - Serial measures of Cardiac autonomic activity Time domain measures

Heart rate increased 48h after surgery in both groups of patients (p<0.001), however there was no difference between PMI groups. (Figure 3) Supine RMSSD values were higher in the PMI group before surgery compared to patients who did not develop PMI [45.7ms (95%CI: 33.5-57.9ms) vs 29.7ms (95%CI: 23.2-36.2ms), p <0.001]. However, patients who developed PMI showed a decrease in RMSSD 24h after surgery [45.7ms (95%CI: 33.5-57.9ms) vs 25.1ms (95%CI: 13.1-37.0) p<0.001], whereas patients who did not develop PMI showed no change [29.7ms (95%CI:23.2-36.2ms) vs 32.7ms (95%CI: 25.9-39.4ms), p=0.521]. A decrease in SDNN 24h after surgery was also only observed in the patients who developed PMI. [36.9ms (95%CI: 28.5-45.4ms) vs 22.5ms (95%CI:14.2-30.8)]. (Supplementary Table 3). The same trend was observed for the recordings in the 45 degree head up position.

### Frequency domain measures

Natural logarithmic High frequency power spectral band (cardiac vagal activity) decreased 24h and 48h after surgery only in patients who developed PMI (p <0.01). (Figure 4). Natural logarithmic low frequency spectral band also decreased after surgery (p<0.001) however this was observed independent of patients having a raised troponin. (Supplementary Figure 2).

**Figure 4:**
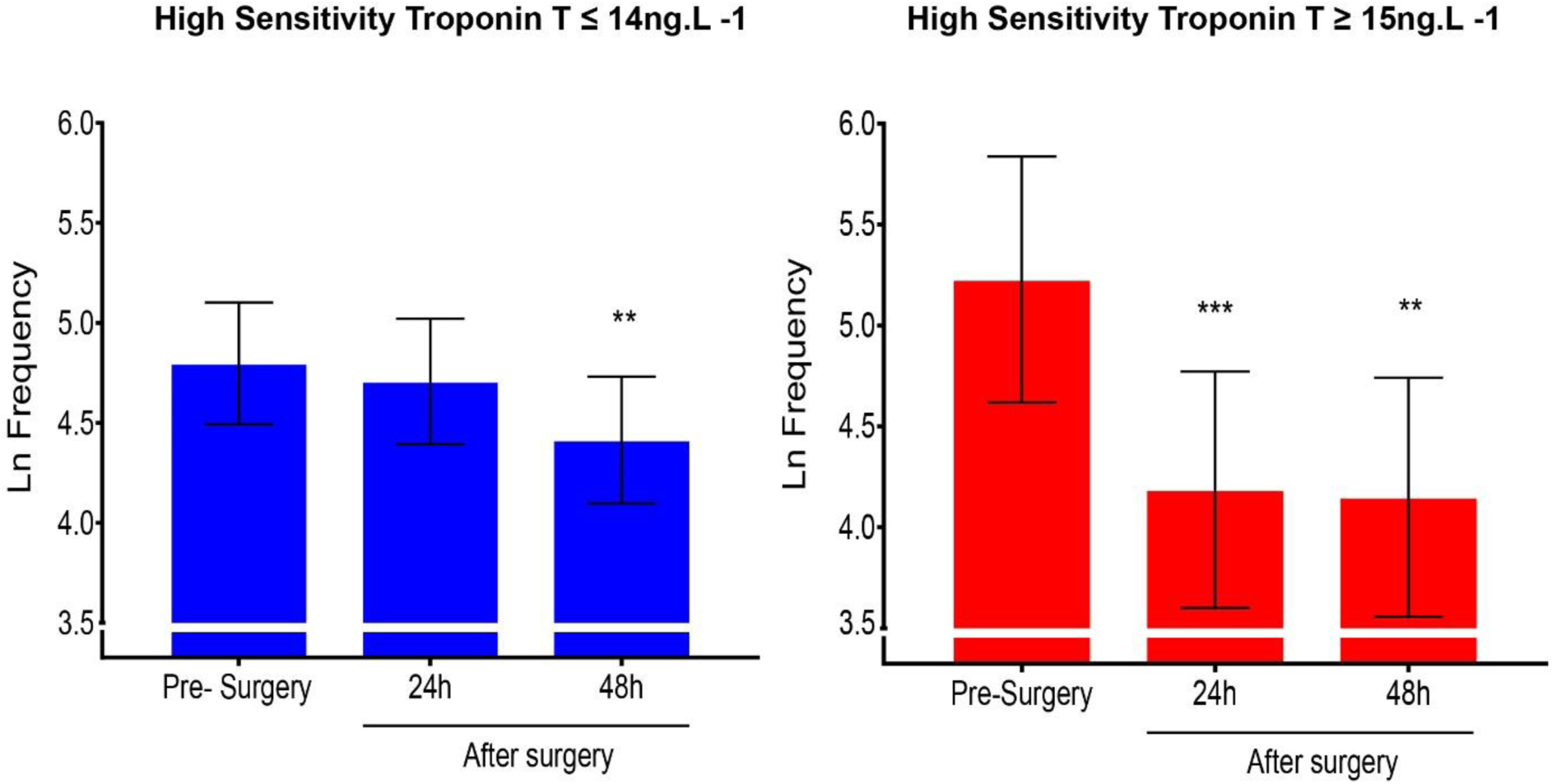
Serial changes in high frequency power spectral analysis (Cardiac parasympathetic activity). Data is presented as natural logarithmic transformed mean values with 95% confidence intervals whilst in the 45 degree head up position. P value refers to day of surgery vs 24h after surgery and day of surgery vs 48h after surgery comparisons (analysed by Mixed model repeated measures, Bonferroni’s test). ** p<0.01, *** p<0.001

**Figure 5:**
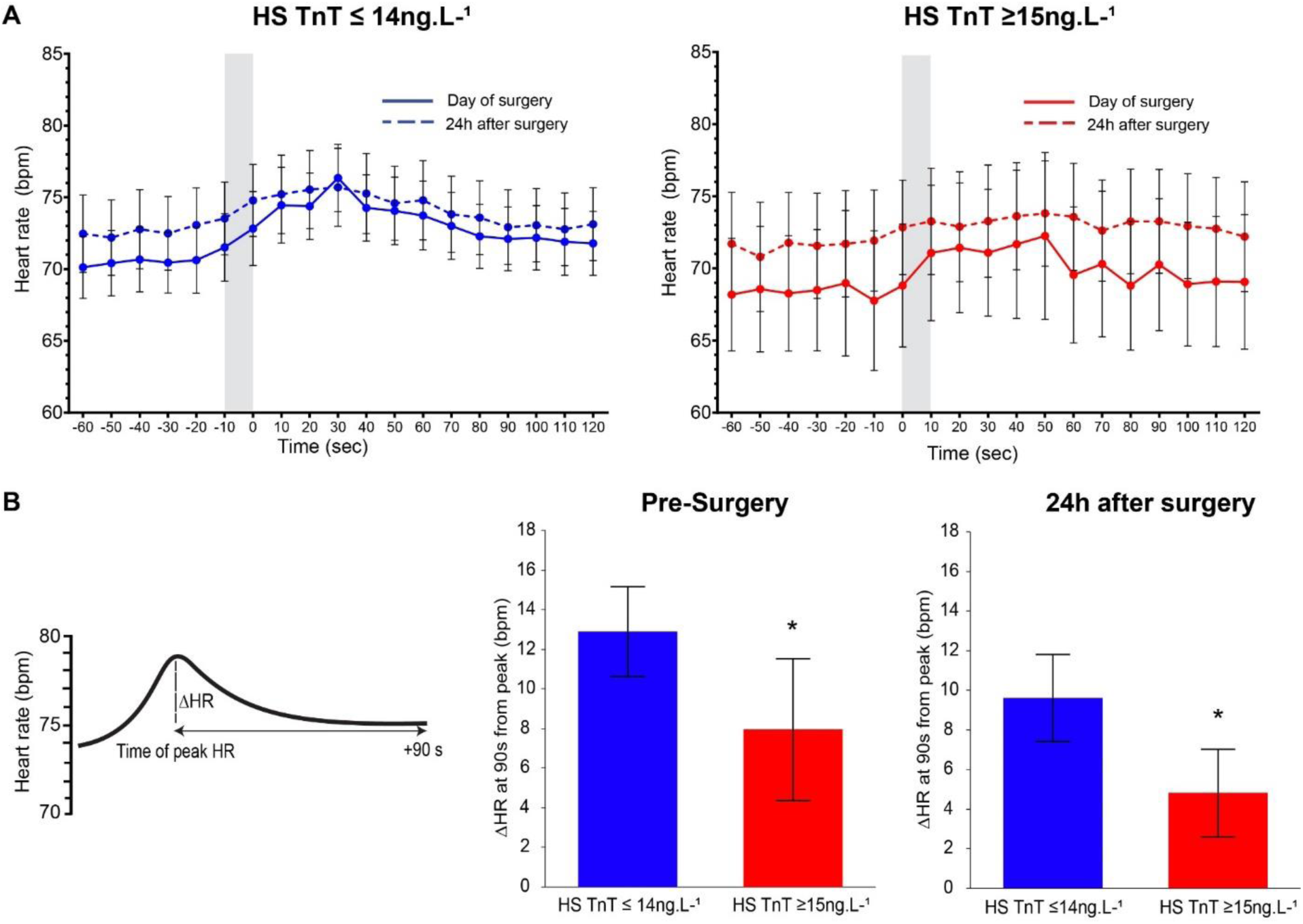
Heart rate changes across standardised orthostatic challenge. **A**: Mean heart rate (95% CI) values before and after the standardised orthostatic challenge (grey shaded area) **B**: Mean Heart rate recovery (95% CI) after 90 s from peak heart rate. P values refers to HsTnT≥15ng.L^−1^ vs HsTnT ≤ 14 ng.L^−1^ (Analysed using one-way ANOVA) * p<0.05.

### Orthostatic challenge-heart rate

Supine heart rate prior to the orthostatic challenge (Figure 4) on the day of surgery and 24h after surgery were similar irrespective if the patients developed PMI (mean difference 2 bpm (95% CI 3-6 bpm, p = 0.394). No differences were observed for when patients achieved their peak heart rate after the orthostatic challenge (PMI group; mean: 30s (20-40s) vs no PMI mean: 30s (20-50s), p = 0.361). Heart rate recovery after a standardised orthostatic challenge 24h after surgery was slower in patients with hsTnT≥15ng.L^−1^ (5 beats minute^−1^ (95%CI: 3 – 7), compared to rapid heart rate recovery in patients who remained who remained free of myocardial injury (10 beats minute^−1^ (95%CI:7 to 12]; p = 0.02). HRR for each 10s interval after peak heart rate is shown in Supplementary Figure 3.

## Discussion

The principle finding of our study is that patients undergoing non-cardiac surgery who developed PMI had a decrease in cardiac parasympathetic activity within 24h of surgery. The loss of cardiac parasympathetic function was demonstrated in both time-domain and frequency domain heart rate variability measures. In addition, delayed heart rate recovery to a standardised orthostatic challenge demonstrated a loss of the cardiac vagal mediated baroreflex response in patients who developed PMI. The cohort of patients in this study were low risk for cardiovascular complications as defined RCRI.

The association of delayed HRR before surgery and PMI after the standardised orthostatic challenge is consistent with a large multicentre cohort study that demonstrated the same association with delayed heart rate recovery, quantified using CPET before noncardiac surgery(4). Using a standardised orthostatic challenge, we were able demonstrate that delayed HRR and its association with PMI persist 24h after surgery.

Frequency domain heart rate variability measures have been shown to be useful screening tools for haemodynamic instability in the perioperative period(18). The impairment of HRR may in part be due to acquired parasympathetic dysfunction as demonstrated by the loss in high frequency power spectrum found in PMI patients with impaired HRR. This supports our previous work that cardiac vagal dysfunction can be identified by baroreceptor insensitivity (15) is associated with an increased post surgery cardiovascular complications(6). The findings that patients who developed PMI were older, male and had a longer duration of surgery was consistent with previous larger observational studies.(19) The higher proportion of active cancer patients in the PMI group of patients is unsurprising considering patients presenting for cancer surgery have impaired cardiovascular and autonomic function (20).

Our study cohort was at low risk for cardiovascular events as assessed by the RCRI. No myocardial infarctions were observed in our study, although we cannot exclude the possibility of silent events. A similar trend of rise in postoperative troponin levels has previously been reported in young health adults after non cardiac surgery, (21) further suggesting non ischaemic mechanisms for PMI.

Reduced cardiac parasympathetic activity contributes to myocardial injury after surgery(22). Translational and experimental in-vivo models have demonstrated that parasympathetic neurotransmitters released by vagal nerve activity confer cardiac organ protection(23), in part through limiting systemic inflammation.(24, 25)

The association between loss of cardiac parasympathetic function and PMI may be explained by the physiological mechanisms through which cardiac vagal function protects the heart. In-vivo experimental models have shown that cardiac vagal function has anti-inflammatory effects (26, 27) which may limit myocardial injury. This anti-inflammatory effect is conferred both directly and via an inflammatory-neural reflex.(25, 28)

### Strengths and Limitations

Our study is the largest observational study measuring serial HRV variables in the perioperative period conducted thus far. It has also demonstrated the feasibility of standardised autonomic orthostatic challenge to assess HRR before and after surgery. Loss of ECG data for HRV analysis was an anticipated factor in this study due to the threshold set for excluding ECG with artefact. However the scale of exclusion was consistent with other HRV and longitudinal studies comparing haemodynamic parametres(8, 17). A further limitation is that the study was not designed to establish benchmark values for time and frequency measures in the perioperative period.

## Conclusions

In summary, these observational data provide further support for the hypothesis that impaired cardiac vagal activity contributes to perioperative myocardial injury. A standardised autonomic orthostatic challenge has potential to assess HRR in the perioperative period.

## Author contributions

GLA, SMM designed the analysis plan. SMM, SK, AR, GM, LGP obtained ECG data. AR, GM, LGP recorded morbidity data. RCM and DB oversaw conduct of study. GLA, SMM performed the data analysis independently. The manuscript was drafted by GLA SMM and revised following critical review by all authors.

## Declaration of competing interests

GLA is an Editor for Intensive Care Medicine Experimental and British Journal of Anaesthesia. GLA has undertaken consultancy work for GlaxoSmithKline; there are no other relationships or activities that could appear to have influenced the submitted work.

## Sources of funding

GLA is supported by British Journal of Anaesthesia/Royal College of Anaesthetists basic science Career Development award, British Oxygen Company research chair grant in anaesthesia from the Royal College of Anaesthetists and British Heart Foundation Programme Grant (RG/14/4/30736).

## References

1. Botto F, Alonso-Coello P, Chan MT, Villar JC, Xavier D, Srinathan S, et al. Myocardial injury after noncardiac surgery: a large, international, prospective cohort study establishing diagnostic criteria, characteristics, predictors, and 30-day outcomes. Anesthesiology. 2014;120(3):564–78.

2. Devereaux PJ, Biccard BM, Sigamani A, Xavier D, Chan MTV, Srinathan SK, et al. Association of Postoperative High-Sensitivity Troponin Levels With Myocardial Injury and 30-Day Mortality Among Patients Undergoing Noncardiac Surgery. JAMA. 2017;317(16):1642-.

3. Cole CR, Blackstone EH, Pashkow FJ, Snader CE, Lauer MS. Heart-rate recovery immediately after exercise as a predictor of mortality. N Engl J Med. 1999;341(18):1351–7.

4. Abbott TEF, Pearse RM, Cuthbertson BH, Wijeysundera DN, Ackland GL, investigators Ms. Cardiac vagal dysfunction and myocardial injury after non-cardiac surgery: a planned secondary analysis of the measurement of Exercise Tolerance before surgery study. Br J Anaesth. 2019;122(2):188–97.

5. Heusser K, Tank J, Luft FC, Jordan J. Baroreflex failure. Hypertension. 2005;45(5):834–9.

6. Toner A, Jenkins N, Ackland GL, Investigators P-OS. Baroreflex impairment and morbidity after major surgery. Br J Anaesth. 2016;117(3):324–31.

7. McCrory C, Berkman LF, Nolan H, O’Leary N, Foley M, Kenny RA. Speed of Heart Rate Recovery in Response to Orthostatic Challenge. Circ Res. 2016;119(5):666–75.

8. Aries MJ, Bakker DC, Stewart RE, De Keyser J, Elting JW, Thien T, et al. Exaggerated postural blood pressure rise is related to a favorable outcome in patients with acute ischemic stroke. Stroke. 2012;43(1):92–6.

9. von Elm E, Altman DG, Egger M, Pocock SJ, Gotzsche PC, Vandenbroucke JP, et al. The Strengthening the Reporting of Observational Studies in Epidemiology (STROBE) statement: guidelines for reporting observational studies. J Clin Epidemiol. 2008;61(4):344–9.

10. Cobas. Troponin T Hs. 2015.

11. Grocott MP, Browne JP, Van der Meulen J, Matejowsky C, Mutch M, Hamilton MA, et al. The Postoperative Morbidity Survey was validated and used to describe morbidity after major surgery. J Clin Epidemiol. 2007;60(9):919–28.

12. Tarvainen MP, Niskanen JP, Lipponen JA, Ranta-Aho PO, Karjalainen PA. Kubios HRV--heart rate variability analysis software. Comput Methods Programs Biomed. 2014;113(1):210–20.

13. Electrophysiology TFotESoCtNASoP. Heart Rate Variability: Standards of Measurement, Physiological Interpretation, and Clinical Use. Circulation Research. 1996(93):1043–65.

14. Goldstein DS, Bentho O, Park MY, Sharabi Y. Low-frequency power of heart rate variability is not a measure of cardiac sympathetic tone but may be a measure of modulation of cardiac autonomic outflows by baroreflexes. Exp Physiol. 2011;96(12):1255–61.

15. Ackland GL, Whittle J, Toner A, Machhada A, Del Arroyo AG, Sciuso A, et al. Molecular Mechanisms Linking Autonomic Dysfunction and Impaired Cardiac Contractility in Critical Illness. Crit Care Med. 2016;44(8):e614–24.

16. Casagrande JT, Pike MC, Smith PG. An Improved Approximate Formula for Calculating Sample Sizes for Comparing Two Binomial Distributions. Biometrics. 1978;34(3).

17. Wieske L, Chan Pin Yin DR, Verhamme C, Schultz MJ, van Schaik IN, Horn J. Autonomic dysfunction in ICU-acquired weakness: a prospective observational pilot study. Intensive Care Med. 2013;39(9):1610–7.

18. Padley JR, Ben-Menachem E. Low pre-operative heart rate variability and complexity are associated with hypotension after anesthesia induction in major abdominal surgery. J Clin Monit Comput. 2018;32(2):245–52.

19. Abbott TE, Ackland GL, Archbold RA, Wragg A, Kam E, Ahmad T, et al. Preoperative heart rate and myocardial injury after non-cardiac surgery: results of a predefined secondary analysis of the VISION study. Br J Anaesth. 2016;117(2):172–81.

20. Cramer L, Hildebrandt B, Kung T, Wichmann K, Springer J, Doehner W, et al. Cardiovascular function and predictors of exercise capacity in patients with colorectal cancer. J Am Coll Cardiol. 2014;64(13):1310–9.

21. Duma A, Wagner C, Titz M, Maleczek M, Hupfl M, Weihs VB, et al. High-sensitivity cardiac troponin T in young, healthy adults undergoing non-cardiac surgery. Br J Anaesth. 2018;120(2):291– 8.

22. David Amar MMF, PhD; Carol B. Pantuck, BA; Harry Shamoon, MD; Hao Zhang, MD; Nancy Roistacher, MD; Denis H. Y. Leung, PhD; Ilana Ginsburg, RN; Richard M. Smiley, MD, PhD. Persistent Alterations of the Autonomic Nervous System after Noncardiac Surgery. Anesthesiology. 1998;89:30–42.

23. Mastitskaya S, Marina N, Gourine A, Gilbey MP, Spyer KM, Teschemacher AG, et al. Cardioprotection evoked by remote ischaemic preconditioning is critically dependent on the activity of vagal pre-ganglionic neurones. Cardiovascular Research. 2012;95(4):487–94.

24. Chavan SS, Pavlov VA, Tracey KJ. Mechanisms and Therapeutic Relevance of Neuro-immune Communication. Immunity. 2017;46(6):927–42.

25. Andersson U, Tracey KJ. Neural reflexes in inflammation and immunity. J Exp Med. 2012;209(6):1057–68.

26. Leib C, Goser S, Luthje D, Ottl R, Tretter T, Lasitschka F, et al. Role of the cholinergic antiinflammatory pathway in murine autoimmune myocarditis. Circ Res. 2011;109(2):130–40.

27. Zhang Y, Popovic ZB, Bibevski S, Fakhry I, Sica DA, Van Wagoner DR, et al. Chronic vagus nerve stimulation improves autonomic control and attenuates systemic inflammation and heart failure progression in a canine high-rate pacing model. Circ Heart Fail. 2009;2(6):692–9.

28. Andersson U, Tracey KJ. Reflex principles of immunological homeostasis. Annu Rev Immunol. 2012;30:313–35.

29. KDIGO 2012 Clinical Practice Guidelinefor the Evaluation and Management ofChronic Kidney Disease. Kidney Int Suppl (2011). 2013;3(1):5–14.

